# DNA aptamers to interfere with HasA, the hemophore of the heme assimilation system in *Pseudomonas aeruginosa,* as a potential antimicrobial strategy

**DOI:** 10.64898/2026.09.03.749098

**Authors:** Federico Bosetto, Tongyuan Wei, Katherine Stott, Stephen H. McLaughlin, Laura S. Itzhaki, Maria Zacharopoulou, Ioanna Mela

## Abstract

*Pseudomonas aeruginosa* relies on heme acquisition to sustain growth and virulence in the iron-limited environment of the host, particularly in the acidic airways of patients with cystic fibrosis where antibiotic efficacy is markedly reduced. The Has heme assimilation system is initiated by the secreted hemophore HasA, which binds extracellular heme and delivers it to the outer-membrane receptor HasR. During infection, HasA is proteolytically processed, generating a truncated form that constitutes the biologically relevant species. Here, we characterise the structural and functional properties of truncated HasA under disease-associated acidic conditions (pH 6.5) and perform a rationally designed Systemic Evolution of Ligands by EXponential enrichment (SELEX) method to identify DNA aptamers capable of binding this hemophore.

Biophysical analyses revealed that truncated HasA is folded, displays a mixed α/β secondary structure, and exists as a concentration-dependent mixture of monomers and domain-swapped dimers. Circular dichroism and nano-differential scanning fluorimetry identified two thermal transitions, consistent with the coexistence of apo/holo and monomer/dimer species. The apo protein bound heme with high affinity (*K*_D_ = 113 nM) and a 1:1 stoichiometry, confirming preservation of its functional binding mechanism. Twelve rounds of SELEX, incorporating target switching and platform switching, yielded a highly enriched aptamer pool dominated by two sequences, HasA_1 and HasA_2. ELONA (Enzyme-Linked Oligonucleotide Assay) assays demonstrated that both aptamers bind specifically to truncated HasA, and biolayer interferometry revealed nanomolar dissociation constants (*K*_D_ = 0.59 µM and 0.32 µM, respectively). These aptamers did not bind full-length HasA, BSA, or control sequences, confirming that selection under acidic conditions drove specificity toward the physiologically relevant form of the hemophore.

Our findings identify HasA-binding DNA aptamers that retain function in acidic environments where antibiotic potency is compromised, highlighting their potential as molecular tools to disrupt heme acquisition in *P. aeruginosa*. This work establishes a foundation for developing aptamer-based antimicrobial strategies targeting the Has system, particularly under acidic conditions where conventional antibiotics are compromised.

## Introduction

*Pseudomonas aeruginosa* is a highly adaptable Gram-negative bacterium capable of thriving in nutrient poor and diverse environmental conditions, enabling its persistence in both community and clinical settings. It is a major opportunistic pathogen in individuals with cystic fibrosis and in immunocompromised patients, where it contributes significantly to morbidity and mortality^1–3^. Its intrinsic, acquired, and adaptive resistance mechanisms confer tolerance to multiple antibiotic classes, including aminoglycosides, quinolones, and β-lactams^1,3–5^. Acidic microenvironments, such as those characteristics of cystic fibrosis lungs, further promote bacterial growth and biofilm formation and markedly reduce the efficacy of several antibiotics^6,7^. As a result, *P. aeruginosa* is listed among the ESKAPE group of multidrug-resistant pathogens, underscoring the urgent need for new therapeutic strategies^8^.

Iron is an essential micronutrient for *P. aeruginosa*, and many other bacteria, supporting key cellular processes, such as energy production, DNA replication, electron transport and virulence^9–11^. In vertebrate hosts most iron is sequestered within heme, a strategy of nutritional immunity that restricts microbial proliferation^12–14^. To overcome this barrier, *P. aeruginosa* employs three heme uptake pathways to obtain iron from the host: the *Has* heme assimilation system^9,10,13,15^, the *Phu* heme uptake system^9,10,15^ and a TonB-dependent mechanism exemplified by HxuC^17^. Among these, the Has system is unique in relying on a secreted hemophore, HasA, which binds extracellular heme and delivers it to the outer-membrane receptor HasR^9,10,15,16^. Because HasA operates at the host–pathogen interface and is required for efficient heme scavenging under iron limitation, it represents an attractive target for antimicrobial intervention.

HasA is secreted into the extracellular space with a C-terminal signal sequence that is subsequently cleaved by *P. aeruginosa* proteases, generating the truncated form that predominates during infection^18^. The final 15–21 C-terminal residues constitute a flexible, intrinsically disordered region reported to interfere with efficient heme uptake, underscoring the biological relevance of the truncated protein^19–21^.

HasA has been extensively characterised in *Serratia marcescens*. Although *P. aeruginosa* HasA shares only ∼50% sequence identity with the *S. marcescens* hemophore^18^, both proteins adopt highly similar tertiary structures and coordinate heme through the same residues^19^. HasA belongs to the αβ protein family and exhibits a structural composition consisting of approximately 30% α-helices and 30% β-strands^22^. Upon heme binding, HasA adopts a distinctive structure featuring adjacent α-helix and β-sheet walls, from which two loops extend to coordinate the heme via His32 and Tyr75^19,22,23^.

HasA binds to heme following a distinct two-step “induced fit” mechanism: In the apo state, the Tyr75 loop captures heme through hydrophobic, π–π stacking, and van der Waals interactions, forming a high-spin Fe(III) intermediate^24,25^. This intermediate then triggers closure of the His32 loop, driven by hydrogen bonding between the heme propionate groups and the loop backbone, ultimately establishing the His32–Fe coordination bond^24,25^. Kinetic analysis shows that Tyr75-mediated capture is rapid, whereas His32 loop closure is comparatively slow^26,27^. The resulting holo complex exhibits exceptionally high affinity, with a reported association constant of 5.3 × 10¹⁰ M⁻¹ ^23^. In *S. marcescens*, HasA can also form a domain-swapped dimer mediated by a conserved hinge loop, which binds two heme molecules and is proposed to function as a heme reservoir^28^.

Aptamers are small single-stranded synthetic nucleic acids that fold into defined three-dimensional structures and bind diverse molecular targets with high affinity and specificity^29–31^. They can recognize metal ions, small molecules, proteins, and whole cells^32^ and are generated from random oligonucleotide pools by an *in vitro* selection process called Systemic Evolution of Ligands by EXponential enrichment (SELEX)^33,34^. Their strong binding properties and broad target range have enabled applications across diagnostics, biotechnology, and therapeutics^35^. In the context of antimicrobial resistance, aptamers represent promising pharmacological tools capable of targeting virulence factors or interfering with essential metabolic pathways^36^.

In this study, we explored the structure, dimerization, and heme-binding properties of the hemophore HasA in its cleaved form under disease-relevant acidic conditions, and using SELEX identified DNA aptamers capable of disrupting the heme assimilation system in *P. aeruginosa.* Our approach, of targeting a secreted hemophore instead of a bacterial membrane or cytoplasmic protein has the potential to reduce selection pressure, and therefore emergence of resistance, while being able to disrupt the bacterium’s ability to source iron from its environment. The aptamers selected in this study retain function in acidic environments where antibiotic potency is compromised, highlighting their potential therapeutic application.

## Results

### Structural and functional characterizations of the Cʹ terminal truncated HasA protein

Full-length and Cʹ terminal truncated HasA were synthesized with an N-terminal 6xHistidine-tag to enable protein purification by affinity chromatography and to facilitate aptamer selection on immobilized targets (Figure 1a). The constructs included all relevant structural features, including the heme-coordination residues, the domain-swapping hinge loop, and the engineered C-terminal deletion. As expected, removal of the final 21 residues resulted in a mobility shift on SDS–PAGE, from 22.8 kDa for the full-length protein to 20.8 kDa for the truncated variant (Figure 1b). Both protein molecular weights were also confirmed by ESI-MS (Supplementary Figure S1a and b). The purified truncated protein displayed a yellowish colour, consistent with partial heme loading arising from *E. coli* metabolism, indicating the presence of both apo and holo species, with the apo state predominating^21,26,27^.

**Figure 1.**
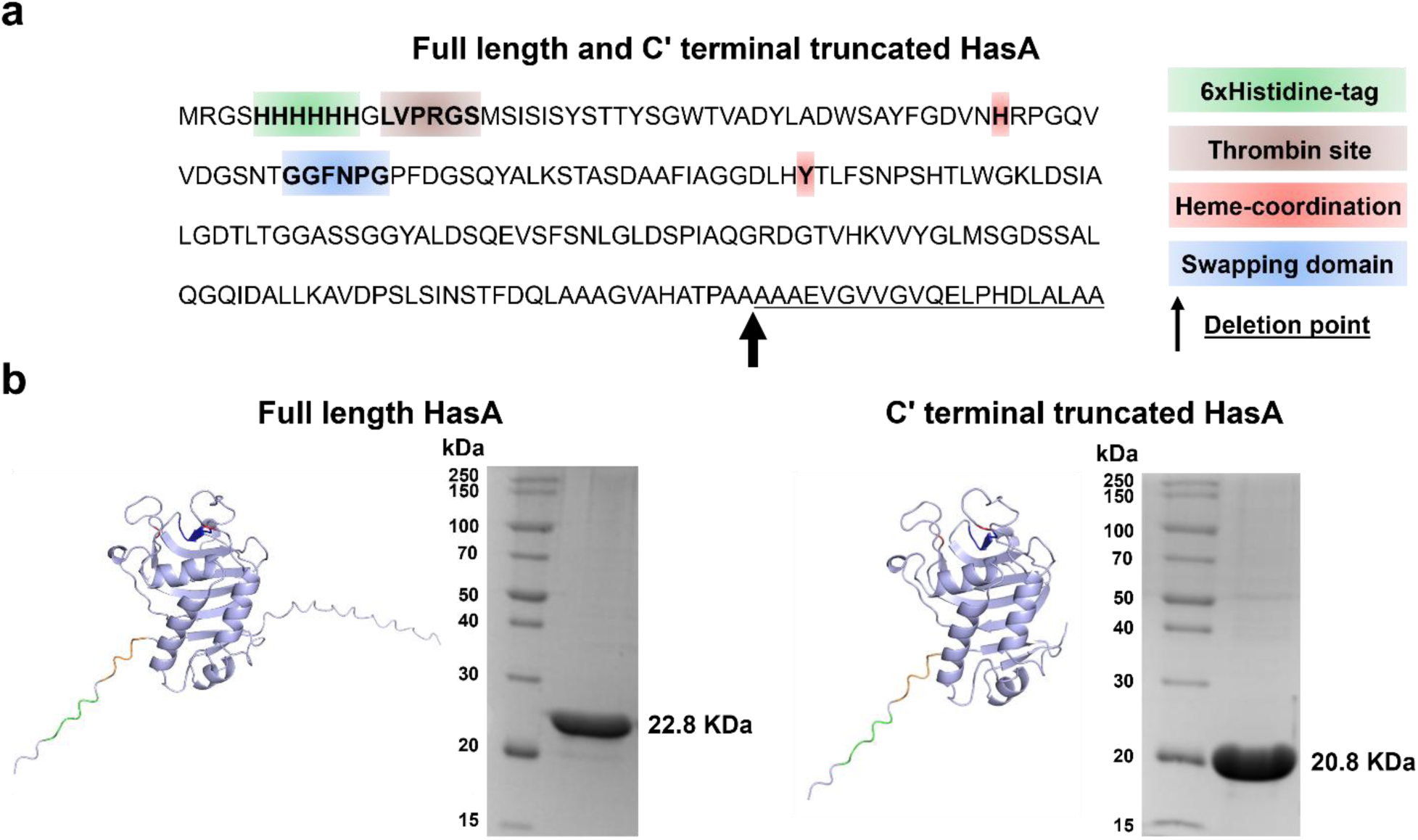
**(a)** Aminoacidic sequences of the full-length and Cʹ terminal truncated HasA proteins, highlighting the 6xHistidine-tag (green), thrombin cleavage site (brown), heme coordination residues (red), swapping domain (blue), and deletion point corresponding to the final 21 Cʹ terminal residues (underlined sequence). **(b)** Both protein variants were successfully purified, and the deletion resulted in a clear mobility shift on SDS-PAGE from 22.8 kDa for the full-length variant to 20.8 kDa for the Cʹ terminal truncated variant.

Circular Dichroism (CD) spectroscopy confirmed that the truncated HasA is folded and exhibits a mixed α/β secondary structure with approximately 26% α-helices (18% regular α-helices and 8% distorted α-helices), 19% antiparallel β-strands (4% left-twisted antiparallel β-strands, 8% relax antiparallel β-strands and 7% right-twisted antiparallel β-strands), 10% parallel β-strands, 10% β-turns and 36% disordered (others) (Figure 2a and b). Thermal denaturation monitored by CD revealed two distinct transitions at ∼49 °C and ∼66 °C, observed at wavelengths characteristic of β-strands (218 nm)^37^ and α-helices (222 nm)^37^ (Figure 2c). NanoDSF (nano-differential scanning fluorimetry) analysis yielded a melting temperature between 62–64 °C across all protein concentrations tested (Figure 2d), consistent with the higher-temperature CD transition. The melting temperature decreases as the protein concentration increases, this could be attributed, on the one hand, to dimer formation and the possible dissociation of the protein and, on the other, to the probable dissociation of the heme group from the holo state.

**Figure 2.**
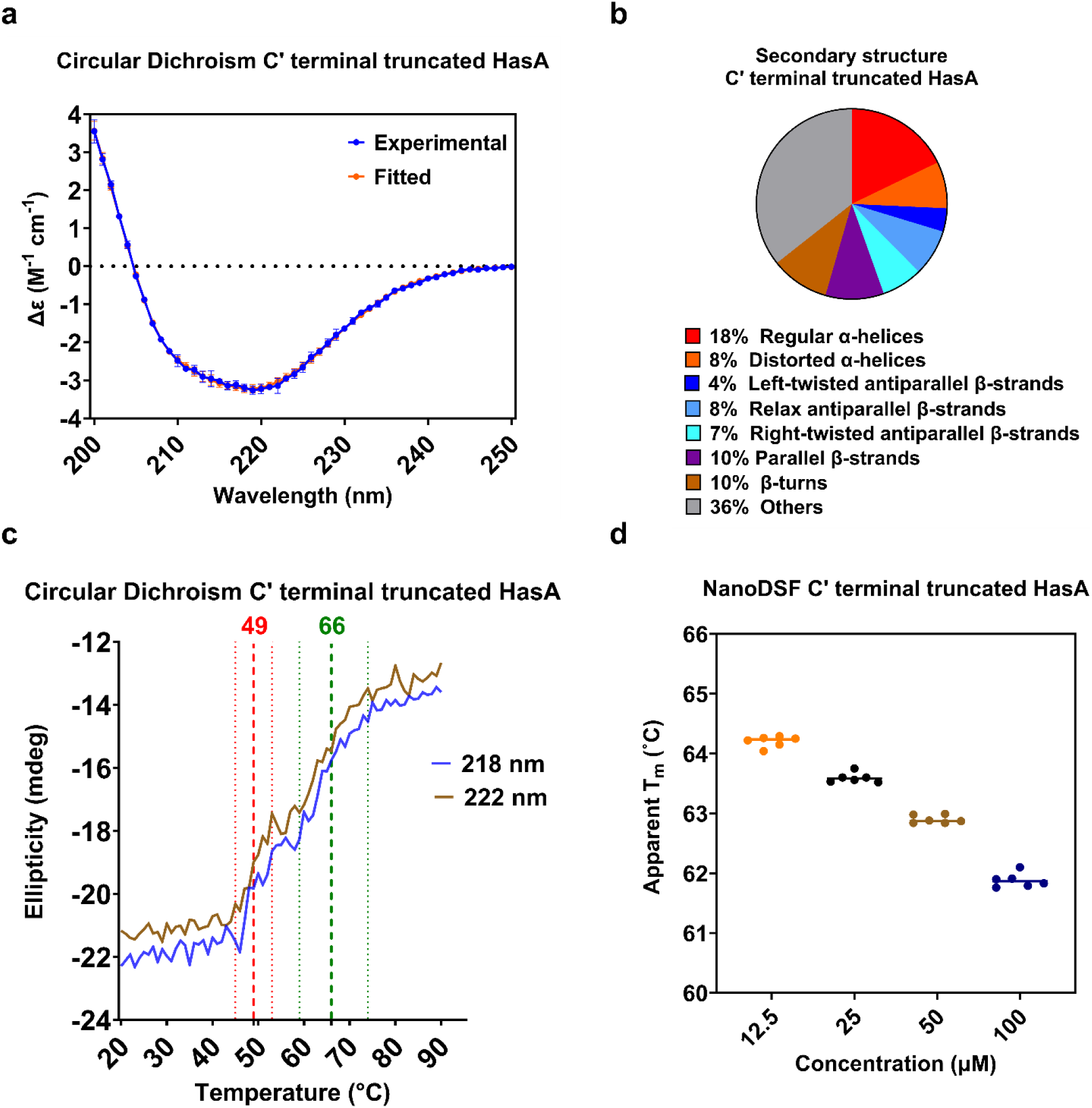
**(a)** CD spectrum and **(b)** resulting secondary structure analysis of the Cʹ terminal truncated HasA using BeStSel indicated a composition of approximately 26% α-helices (18% regular α-helices and 8% distorted α-helices), 19% antiparallel β-strands (4% left-twisted antiparallel β-strands, 8% relax antiparallel β-strands and 7% right-twisted antiparallel β-strands), 10% parallel β-strands, 10% β-turns and 36% disordered (others). **(c)** Thermal denaturation monitored by CD showed two transitions at 49 °C (red dotted line) and 66 °C (green dotted line), as indicated by the two plots measured at 218 nm for β-strands (blue trace) and 222 nm for α-helices (brown trace). **(d)** NanoDSF analysis of the Cʹ terminal truncated HasA revealed a melting temperature between 62 °C and 64 °C across the protein concentrations tested: 12.5 μM (orange), 25 μM (black), 50 μM (brown) and 100 μM (blue).

To assess oligomeric state, sedimentation velocity analytical ultracentrifugation (AUC) was performed on the truncated protein. The interferometric scans revealed a concentration-dependent mixture of monomeric and dimeric species, with peaks at ∼20.0 kDa and ∼36.9 kDa (Figure 3a). The presence of the conserved GGFNPG hinge loop (Figure 1a) suggests that dimerization may occur through a domain-swapping mechanism analogous to that described for *Serratia marcescens* HasA^28^. The coexistence of monomer/dimer and apo/holo species may contribute to the two thermal transitions observed by CD. In particular, the lower-temperature transition (∼49 °C) could reflect unfolding of the monomeric apo state, whereas the higher-temperature transition (∼66 °C) likely corresponds to rearrangements associated with dimer dissociation and/or unfolding of the monomeric holo species stabilized by heme.

**Figure 3.**
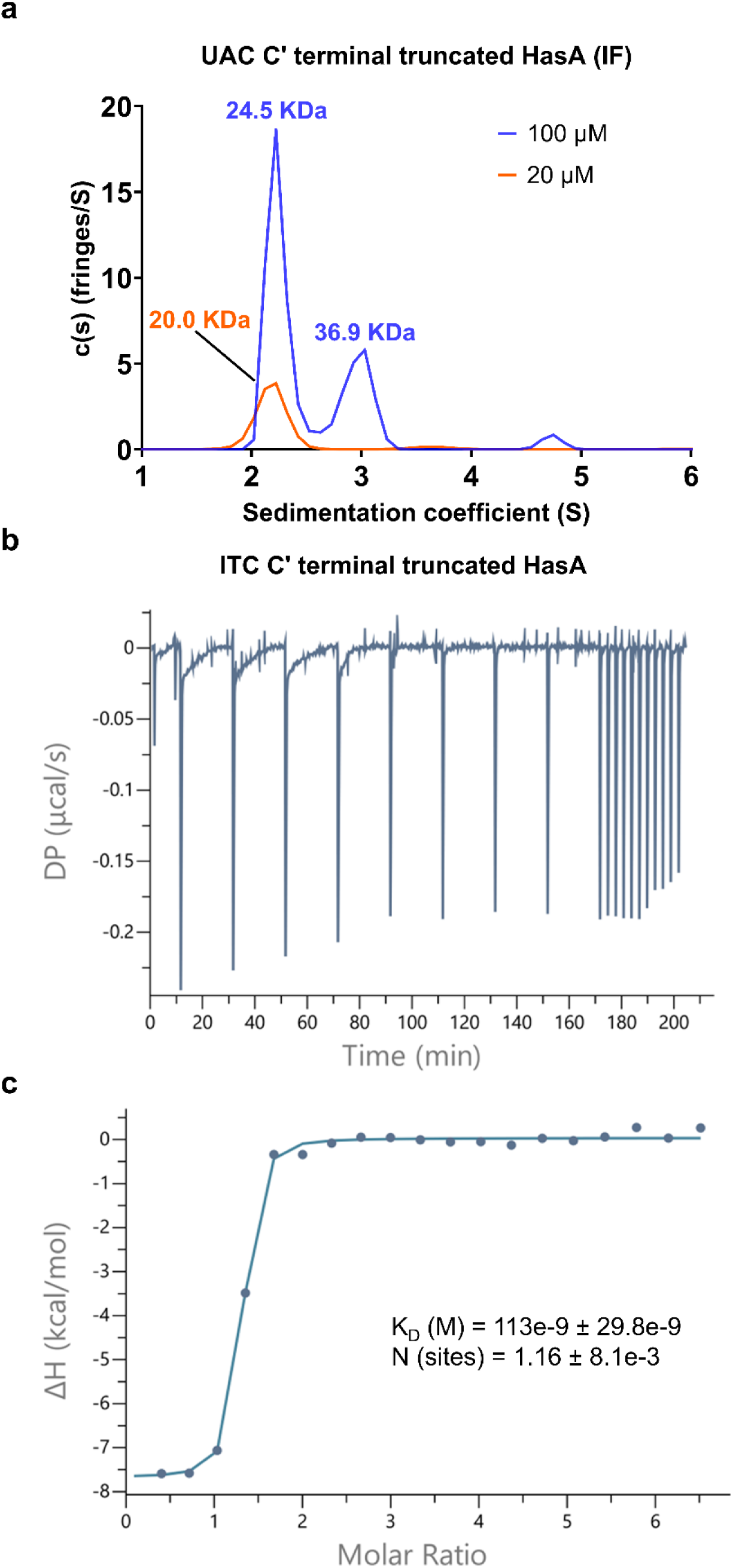
**(a)** AUC of the protein reported a mixed population of monomers and dimers in the interferometer scan (IF): the 20 μM sample showed a single 20.0 kDa peak (orange trace) corresponding to the monomer, whereas the 100 μM sample displayed two peaks at 24.5 kDa and 36.9 kDa (blue trace), indicating dimer formation. ITC of the Cʹ terminal truncated HasA in the presence of heme, showing **(b)** the Differential Power (DP) and **(c)** the integrated and normalised heats per injection.

Heme binding to the apo form of truncated HasA was quantified by isothermal titration calorimetry (ITC). The titration yielded a dissociation constant of 113±30 nM and a stoichiometry of 1.16±0.01, consistent with high-affinity binding and a 1:1 protein-to-heme ratio (Figure 3b and c; Supplementary Table S1). SEC-MALS analysis further confirmed heme association, showing an increase in molecular mass from 23 kDa (apo) to 24.3 kDa (holo), consistent with formation of the heme-bound state (Supplementary Figure S2).

### A DNA aptamer pool targeting the hemophore HasA

The DNA aptamer selection against the hemophore HasA from *P. aeruginosa* was carried out over twelve rounds, initially using the full-length protein (R1–R6) and subsequently the biologically relevant C-terminal truncated form (R7–R12) (Figure 4a; Supplementary Table S2). Because HasA is cleaved extracellularly by *P. aeruginosa* proteases and the truncated form predominates in the infection environment^18^, switching to the truncated protein ensured that selection pressure was applied to the physiologically relevant target. Two immobilization platforms—Ni-NTA magnetic beads and MaxiSorp plates—were used sequentially to refine specificity and broaden epitope accessibility (Figure 4a).

**Figure 4.**
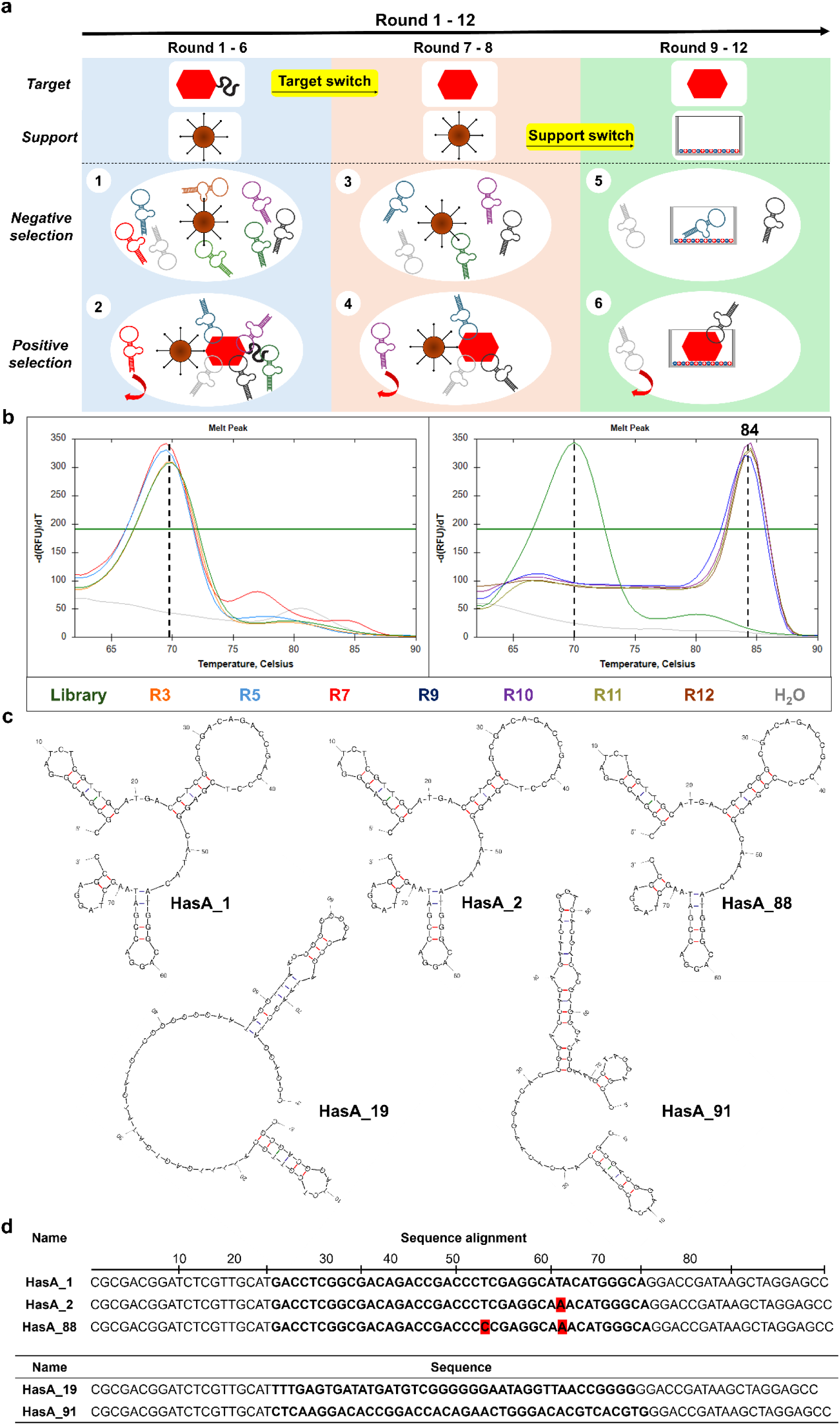
**(a)** Chronological workflow of the 12-round SELEX procedure. The selection strategy was logically partitioned into three sequential phases based on targeted structural domains and supporting matrices. Each round included negative selection on empty beads 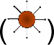 in steps 1 and 3 and on empty MaxiSorp wells 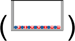 in step 5, and a wash step (red arrow) of non-specifically bound aptamers in steps 2, 4 and 6. Round 1–6, steps 1 and 2: pre-enrichment of the aptamer pool on magnetic beads against the full length protein containing both the structured core 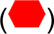 and the unstructured flexible domain 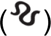. Round 7–8, steps 3 and 4: target-switching phase on magnetic beads, where the selection pressure shifted exclusively to the truncated core domain 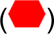, discarding flexible-tail binders. Round 9–12, steps 5 and 6: platform-switching phase on MaxiSorp plate, where the aptamer pool was screened on the immobilized truncated core domain 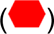. **(b)** Monitoring of the SELEX progression by qPCR and melting curve analysis at R3, R5, R7, R9, R10, R11 and R12. At R3 (orange curve) and R5 (cyan curve) the melting peaks overlap with the one of the DNA library (green curve) at 70 °C, suggesting a high variability of the aptamer pools. Starting from R7 (red curve), the aptamer pool showed convergence toward a set of similar sequences, as evidenced by the appearance of two additional melting peaks at higher temperatures. A complete shift of the melting temperature from 70 °C, characteristic of the DNA library, to 84 °C was visible starting from R9 (blue curve); and for R10 (violet curve), R11 (olive curve) and R12 (brown curve) the melt peak started to narrow, reflecting the selection of an increasingly homogeneous group of sequences. **(c)** Predicted secondary structures for the sequenced aptamers at R12 were obtained by Mfold server at 25 °C and ionic conditions of 150 mM NaCl and 2 mM MgCl_2_. Aptamers HasA_1, HasA_2 and HasA_88 shared the same secondary structure, while aptamers HasA_19 and HasA_91 exhibit different secondary structures. **(d)** Single nucleotide modifications at random regions for aptamers HasA_1, HasA_2 and HasA_88 are reported (red highlight); sequences of HasA_19 and HasA_91 are also listed.

Progression of the SELEX was monitored by qPCR melt-curve analysis, which revealed a clear shift in melting behaviour from 70 °C (characteristic of the starting library) to 84 °C as selection advanced (Figure 4b). Early rounds (R3, R5) showed melting profiles overlapping with the naïve library, indicating high sequence diversity and limited enrichment. Beginning at R7, following the transition to the truncated HasA target, additional higher-temperature melting peaks emerged, suggesting convergence toward a subset of related sequences. From R9 onward, the melt curves displayed a single, sharp peak at 84 °C, consistent with substantial reduction in pool heterogeneity and enrichment of a dominant aptamer population (Figure 4b, right panel). This second transition coincided with the switch to MaxiSorp plate immobilization, where the protein is presented in a non-oriented manner, likely exposing a broader range of epitopes and enabling more stringent selection.

By R10–R12, the melting peak further narrowed, indicating the emergence of a highly homogeneous pool. Next-generation sequencing (NGS) of the R12 pool confirmed this convergence: two aptamers, HasA_1 and HasA_2, accounted for ∼82% of all sequences (75% and 7%, respectively) (Supplementary Table S3). Secondary-structure predictions showed that HasA_1 and HasA_2 adopt the same fold as the HasA_88 aptamer, which was also examined experimentally (Figure 4c and d). Although HasA_19 and HasA_91 were only minimally enriched, they were retained for further analysis because their predicted secondary structures differed from the dominant motif, providing structural diversity within the candidate set (Figure 4c and d).

### HasA_1 and HasA_2 aptamers bind specifically to Cʹ terminal truncated HasA

Binding of the selected DNA aptamers to HasA was first assessed using an ELONA assay coupled to a qPCR read-out, enabling quantitative comparison of aptamer interactions with full-length HasA, Cʹ terminal truncated HasA, and BSA as a negative control (Figure 5a; Supplementary Table S4). Among the candidates, HasA_1 and HasA_2 displayed clear and specific affinity for the truncated protein. HasA_1 generated a signal approximately seven-fold higher than BSA (7.70 × 10^-5^ pmol vs. 1.05 × 10^-5^ pmol), while HasA_2 produced a five-fold increase (5.28 × 10^-5^ pmol vs. 1.07 × 10^-5^ pmol). HasA_1 also bound the truncated protein ∼3.5-fold more strongly than the naïve DNA library (2.20 × 10^-5^ pmol) and exhibited roughly two-fold higher binding than to full-length HasA (3.88 × 10^-5^ pmol). In contrast, HasA_19, HasA_91, and HasA_88 showed no significant binding relative to BSA or the DNA library. These results identify HasA_1 and HasA_2 as the most promising aptamers for specific recognition of the biologically relevant truncated HasA.

**Figure 5.**
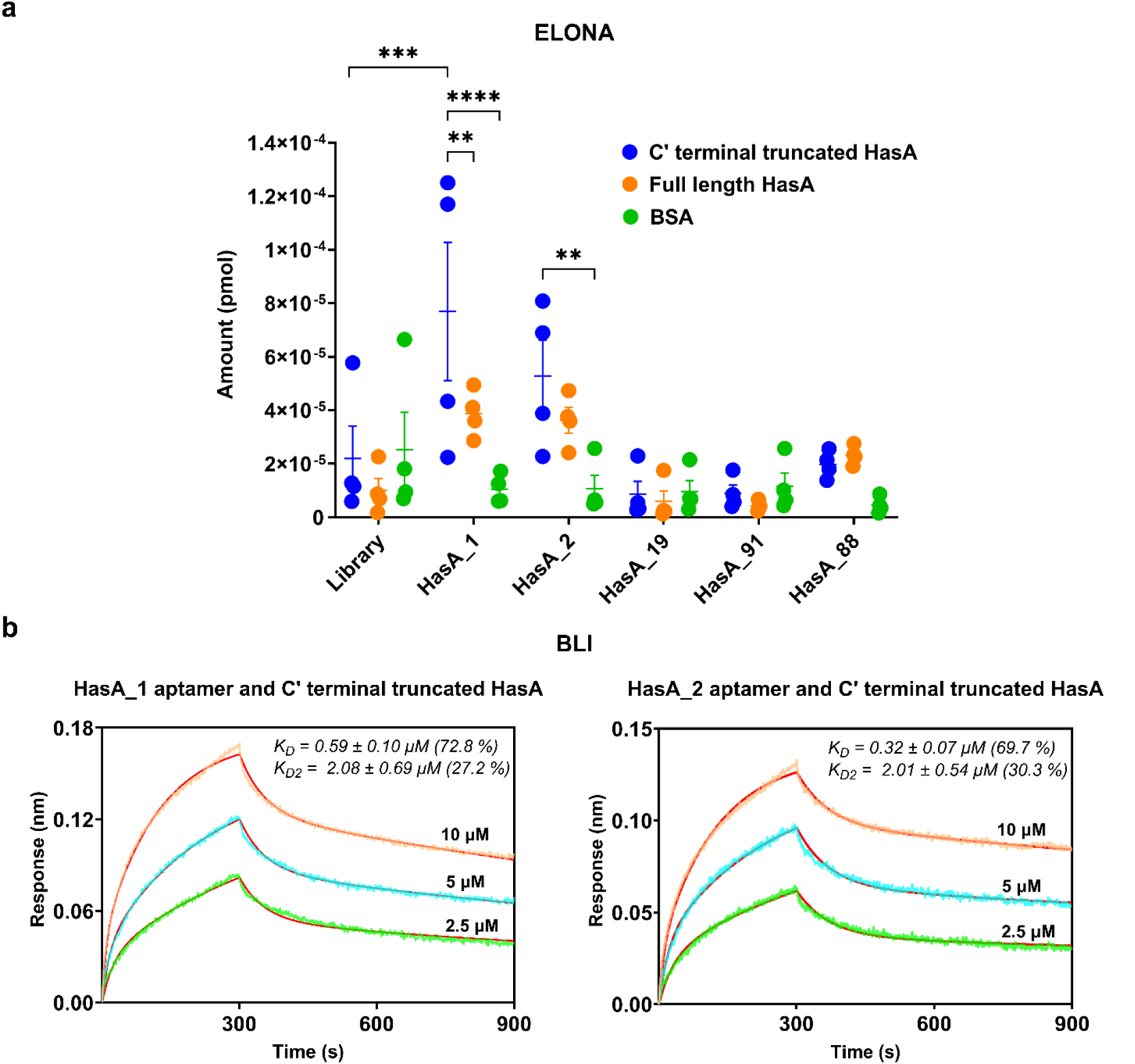
**(a)** ELONA assay coupled with a qPCR read-out was performed in four biological replicates. The assay demonstrated the specificity of both HasA_1 and HasA_2 aptamers to C-terminal truncated HasA, statistical differences compared the binding to BSA were reported (HasA_1 ****P<0.0001, HasA_2 **P=0.0024). HasA_1 aptamer binding to the Cʹ terminal truncated HasA was statistically higher than that of the DNA library (***P=0.0002) and its binding specificity for the truncated form was also statistically relevant when compared with the full length HasA (**P=0.0062). Analysis of the statistical significance was carried out by 2-way ANOVA with Dunnett’s posthoc test for multiple comparisons and with BSA, full length HasA and DNA library used as controls. **(b)** BLI was performed in three biological replicates, aptamer binding curves were obtained using Cʹ terminal truncated HasA protein at concentrations of 10, 5, and 2.5 μM (orange, cyan and green curves, respectively).

Biolayer interferometry (BLI) was then used to characterize the kinetics of aptamer–protein interactions and determine dissociation constants (Figure 5b; Supplementary Table S5). Consistent with the ELONA results, both HasA_1 and HasA_2 bound specifically to the truncated HasA, whereas no measurable binding was observed for the DNA library, the scramble sequence (Supplementary Figure S4), or the remaining aptamer candidates (HasA_19, HasA_91, HasA_88). The sensorgrams for HasA_1 and HasA_2 exhibited complex binding behaviour, likely reflecting interaction with a heterogeneous protein population comprising monomeric and dimeric species as well as apo and holo forms. Accordingly, the data were best fitted using a 2:1 heterogeneous binding model. This analysis yielded *K*_D_ values of 0.59±0.10 µM (HasA_1) and 0.32±0.07 µM (HasA_2), together with secondary *K*_D2_ values of 2.08±0.69 µM and 2.01±0.54 µM, respectively (Figure 5b; Supplementary Table S5).

Together, the ELONA and BLI results demonstrate that HasA_1 and HasA_2 specifically recognize the Cʹ terminal truncated form of HasA and form measurable complexes with nanomolar affinity, validating them as lead aptamer candidates for targeting the Has heme assimilation pathway.

## Discussion

*Pseudomonas aeruginosa* remains a major clinical challenge in the context of antimicrobial resistance, particularly in the acidic microenvironment of cystic fibrosis lungs, where conventional antibiotics lose potency and the bacterium displays enhanced tolerance^6,7^. Developing therapeutic strategies that target essential virulence pathways rather than relying solely on bactericidal mechanisms is therefore an increasingly important goal. In this study, we focused on the hemophore HasA, a key component of the Has heme assimilation system, with the aim of identifying DNA aptamers capable of binding the biologically relevant truncated form of the protein under disease-associated acidic conditions.

We first characterized the Cʹ terminal truncated HasA, which represents the predominant extracellular form generated by proteolytic cleavage in *P. aeruginosa*^18^. CD analysis confirmed that the truncated protein is folded and displays a mixed α/β secondary structure. Interestingly, the α-helical and β-strand content showed consistency with previously reported values for full-length HasA^22^, despite differences regarding the source organism, the truncated form, the analysis conditions (pH 6.5), and the likely different methods used to estimate the secondary structure content. Thermal denaturation revealed two distinct transitions, and NanoDSF supported the higher-temperature unfolding event. Analytical ultracentrifugation demonstrated that truncated HasA exists as a mixture of monomers and dimers, consistent with the domain-swapped dimerization described for *Serratia marcescens* HasA^28^. The coexistence of monomeric/dimeric and apo/holo species may contribute to the biphasic thermal behaviour observed by CD.

Heme titration by ITC showed that the apo form of truncated HasA binds heme with high affinity (*K*_D_ = 113 nM) and a stoichiometry of 1:1, in agreement with previous studies^38^. SEC-MALS further supported heme association, revealing a modest increase in molecular mass upon ligand binding. Together, these results establish that the truncated protein retains the structural and functional properties required for heme acquisition and is therefore an appropriate target for aptamer selection.

The SELEX strategy was designed to progressively refine specificity toward the truncated HasA. Early rounds against full-length HasA maintained high sequence diversity, whereas switching to the truncated protein at R7 produced a clear shift in melting behaviour, indicating enrichment toward a subset of sequences. A second transition occurred upon moving to MaxiSorp plate immobilization, likely due to improved epitope accessibility. By R9–R12, the aptamer pool had converged to a highly homogeneous population, and NGS analysis revealed that two aptamers—HasA_1 and HasA_2—dominated the pool, together accounting for ∼82% of all sequences.

Binding assays confirmed the specificity of these aptamers. ELONA demonstrated that HasA_1 and HasA_2 bind the truncated protein with significantly higher signals than BSA, the DNA library, or full-length HasA. BLI further validated these interactions and provided kinetic parameters, with nanomolar dissociation constants for both aptamers. The heterogeneous binding behaviour observed in BLI likely reflects interaction with a mixed population of HasA species (monomer/dimer and apo/holo), consistent with our biophysical characterization. Although the affinities of HasA_1 and HasA_2 are modest relative to the extremely high affinity of HasA for heme, they nonetheless represent promising lead candidates for targeting the Has system. Future optimisation—such as multivalent formats or chemical modifications—may further enhance their potency.

## Conclusions and future perspectives

Overall, we report the detailed biochemical and biophysical characterisation of truncated HasA under physiologically relevant acidic conditions and identify two DNA aptamers capable of binding this hemophore with specificity. These aptamers offer a potential route to interfering with heme acquisition in *P. aeruginosa*, a strategy that could impose metabolic disadvantage under conditions where antibiotic efficacy is compromised. Future studies will be essential to determine how aptamer binding affects heme uptake, whether it impacts other iron-acquisition pathways, and how these effects manifest in *P. aeruginosa* physiology *in vitro*, *ex vivo*, or in cystic fibrosis-associated environments. In parallel, efforts to develop aptamers targeting the outer-membrane receptor HasR may enable comprehensive blockade of the Has system and further advance aptamer-based antimicrobial approaches.

## Acknowledgements

The authors would like to thank Dr Pamela J. E. Rowling and Khushboo Matwani for helpful discussions.

I.M. acknowledges funding from the Royal Society (URF/R1/221795, IES∖R3∖223128, IES∖R2∖222107, RGS∖R1∖231266), the National Biofilms Innovation Centre (BB/R012415/1 03PoC20-105) and the David James Trust.

M.Z. acknowledges funding from Oppenheimer Fellowship (School of the Physical Sciences, University of Cambridge).

## Materials and Methods

### Cloning, expression and purification of full length and Cʹ terminal truncated HasA

The gene encoding for *P. aeruginosa* HasA was obtained from the Pseudomonas Genome Database^39^. The gene sequence was designed with the *BamHI* restriction site at the 5ʹ position and the *HindIII* restriction site at the 3ʹ position. The HasA gene was synthesized from IDT and codon optimized through GeneArt (Thermofisher Scientific) for *E. coli* cells. The gene was also amplified by PCR with Hot Start *Taq* DNA polymerase (New England Biolab, M0495L) and with following forward (5ʹ-TATATGATCACTTGTCTGGTTCCGCGTG-3ʹ) and reverse (5ʹ-TAATAAGCTTTTATGCTGCAGGGGTTGC-3ʹ) primers at 0.1 µM to introduce a deletion of the last 21 amino acids at the Cʹ terminal side of the protein (Figure 1a and b). The amplification protocol was 95 °C for 30 sec, (95 °C for 30 sec, 53 °C for 30 sec and 68 °C for 1 min) x 30 cycles and 68 °C for 5 min. Both gene and PCR product were cloned within pBRE vector harboring a 6-Histidine tag at the 5ʹ position (Figure 1a). Chemically competent Lucigen C41(DE3) *E. coli* cells were transformed with both ligation products and inoculated in 2xYT containing 100 µg/ml ampicillin, then grown overnight at 37 °C and 220 rpm. Overnight cultures were used to inoculate the main cultures with a 1:100 ratio and grown at 37 °C and 220 rpm until an OD_600_ of 0.8-1.0 was measured. Protein expression was induced by the addition of isopropyl-b-D-1-thiogalactopyranoside (IPTG) to a final concentration of 1 mM. Both full length and Cʹ terminal truncated proteins were purified by affinity chromatography with HisTrap^TM^ excel 1 ml column with a flow rate of 1 ml/min using an AKTA Pure system. After sample application, the column was washed with 20 CV in Buffer A2 (50 mM sodium phosphate pH 7.5, 500 mM NaCl) and the elution was performed with 40 CV of a linear gradient of the imidazole of the Buffer B (50 mM sodium phosphate pH 7.5, 150 mM NaCl, 500 mM imidazole) from 0% to 100% followed by 10 CV at 100% Buffer B. Protein aliquots corresponding to the elution peak were recovered, assessed using 15% SDS–PAGE, and dialyzed against 1 liter Buffer A (50 mM sodium phosphate pH 7.5, 150 mM NaCl) at 4 °C, overnight. Protein concentration was determined by UV adsorption set at 280 nm and adjusted with the predicted molar extinction coefficient calculated by ProtParam tool at ExPASy web portal (ɛ= 28420 M^-1^ cm^-1^).

### Electrospray ionisation mass spectrometry (ESI-MS)

ESI-MS was performed on a Xevo G2 mass spectrometer with data analysed using MassLynx software (Waters UK) (Yusuf Hamied Department of Chemistry, University of Cambridge, UK).

### Circular Dichroism (CD)

CD was measured on Chirascan CD spectrometer (Applied Photophysics, Leatherhead, UK) in 1-mm-pathlength Precision Cells (110-QS, Hellma Analytics, Müllheim, Germany), using Cʹ terminal truncated HasA at 10 µM in 50 mM sodium phosphate pH 6.5, 150 mM NaCl buffer. The CD spectrum was recorded in the range of 200-250 nm, at 25 °C, in three scans, and with a bandwidth of 1 nm; the average ellipticity from three biological replicates was reported. CD spectrum was also recorded at 222 nm and at 218 nm, from 20 °C to 90 °C, in one scan, with a step size of 1 °C/min, and bandwidth of 1 nm; the average ellipticity from three biological replicates was reported. Raw CD data were corrected by subtracting the spectrum of the buffer solution. Beta Structure Selection (BeStSel) server was used to determine the protein secondary structure and fold recognition from the CD spectrum, giving an indication of α-helix and β-sheet content^40^.

### Nano Differential Scanning Fluorimetry (NanoDSF)

NanoDSF was performed with a Prometheus NanoDSF instrument (NanoTemper Technologies) to evaluate the Cʹ terminal truncated HasA melting temperature at 12.5, 25, 50 and 100 µM in 50 mM sodium phosphate pH 6.5, 150 mM NaCl buffer. Protein thermal stability was estimated by direct fit of the fluorescence intensity read at 350 nm from 20 °C to 90 °C with a 1 °C/min rate.

### Analytical ultracentrifugation (AUC)

Sedimentation velocity experiments were conducted with an Optima AUC XL-I ultracentrifuge (Beckman). Standard double-sector Epon centrepieces equipped with sapphire windows contained 400 μL of Cʹ terminal truncated HasA at 20 µM and 100 µM in 50 mM sodium phosphate pH 6.5, 150 mM NaCl buffer. Interference data were collected at 50,000 rpm, at 20 °C. Multi-component sedimentation coefficient distributions were obtained using Sedfit v.14.1^41^.

### Isothermal titration calorimetry (ITC)

Isothermal titration calorimetry binding studies were performed on Auto-iTC200 instrument (Malvern Instruments). Hemin (Sigma-Aldrich, 51280) at 600 µM was titrated into the ITC cell containing 19 µM HasA in a series of 19 injections of 2 µL, preceded by a 0.5 µL pre-injection accounting for potential exchange of materials during the equilibration phase of the measurement. The pre-injection heat was not used during fitting. Integrated excess heats from each injection fitted to a single-site binding model using the MicroCal PEAQ-ITC Analysis Software 1.0.0.1258 (Malvern Instruments).

### Size-Exclusion Chromatography with Multi-Angle Light Scattering (SEC-MALS)

SEC-MALS analysis was performed for apo Cʹ terminal truncated HasA, pre SEC - holo state Cʹ terminal truncated HasA and post ITC - holo state Cʹ terminal truncated HasA using an online Dawn Helios ii system (Wyatt) equipped with a QELS+ module (Wyatt) and an Optilab rEX differential refractive index detector (Wyatt). One hundred uL of: apo Cʹ terminal truncated HasA at ∼0.4 mg/ml, pre SEC - holo state Cʹ terminal truncated HasA at ∼0.7 mg/ml, and post ITC - holo state Cʹ terminal truncated HasA, were auto-injected onto a S75 Sephadex 75 HR10/300 Increase column (Cytiva) and run at 0.5 ml/min. The system was equilibrated with SEC buffer (50 mM Sodium phosphate pH 6.5, 150 mM NaCl). The light scattering and protein concentration at each point across the peaks in the chromatograph were used to determine the absolute molecular mass from the intercept of the Debye plot using Zimm’s model as implemented in the ASTRA 7.3.0.11 (Wyatt). To determine inter-detector delay volumes, band-broadening constants and detector intensity normalization constants for the instrument, Bovine Serum Albumin (ThermoFisher Scientific, 23209) was used as a standard prior-to sample measurement. Data was plotted using GraphPad Prism 11.0.2 (GraphPad Software).

### SELEX on magnetic beads

Dynabeads™ magnetic beads (Invitrogen, 10103D) were used to perform SELEX rounds from 1 to 6 against full length HasA and rounds 7 and 8 against Cʹ terminal truncated HasA, as reported in the Supplementary Table S2. The magnetic beads were first washed twice with 100 µl of SBB (50 mM sodium phosphate pH 6.5, 150 mM NaCl) and then bound with the protein of interest for 30 min at room temperature with shaking. The beads-protein complex was washed with 100 µl of SBB five times. DNA library (5ʹ-CGCGACGGATCTCGTTGCAT*N_40_*GGACCGATAAGCTAGGAGCC-3ʹ) was heated up to 95 °C for 8 min in SBB, 2 mM MgCl_2_ was added to the hot solution and immediately transferred to 4 °C for 10 min and then at room temperature for 10 min. Folded DNA library was incubated first with empty magnetic beads as counter selection step and then with the beads-protein complex, as indicated in the Supplementary Table S2. Washing step was performed with 200 µl of SWB (50 mM sodium phosphate pH 6.5, 150 mM NaCl, 2 mM MgCl_2_), the modality of the wash and the SWB composition vary according to the applied stringency of the washing conditions (Supplementary Table S2). DNA aptamers were eluted from beads-protein complex in 40 µl of nuclease-free H_2_O by heating at 95 °C, twice. PCR amplification of the recovered aptamer pool was performed with Hot Start *Taq* DNA polymerase and with the following forward (5ʹ-CGCGACGGATCTCGTTGCAT-3ʹ) and reverse (5ʹ-[Biotin]-GGCTCCTAGCTTATCGGTCC-3ʹ) primers at 0.1 µM. The amplification protocol was 95 °C for 1 min, (95 °C for 15 sec, 60 °C for 15 sec and 68 °C for 20 sec) x N cycles and 68 °C for 5 min. For each selection round the best number of amplification cycle (N) was determined to generate sufficient quantities of the aptamer pool for the consecutive round without introducing PCR artefacts. Streptavidin magnetic beads (New England Biolab, S1420S) were used to regenerate the ssDNA conformation of the aptamer pool after PCR amplification. One mg of streptavidin magnetic beads was washed twice with 1 ml of TEN buffer (10 mM Tris-HCl pH 8, 1 mM EDTA, 1 M NaCl) and incubated with the PCR product, previously diluted in TEN buffer to 1 ml, for 1 hour at room temperature in a rotor. The PCR product-streptavidin magnetic beads complex was washed three times with TEN buffer, and the forward strand was eluted from the beads by alkaline denaturation with 20 mM NaOH for 10 min. The eluted ssDNA was buffer-exchanged to nuclease-free H_2_O by amicon ultra filter MWCO 3 kDa and quantified by UV adsorption set at 260 nm.

### SELEX on plate

Nunc MaxiSorp plate (ThermoFisher Scientific, 439454) was used to perform SELEX rounds from 9 to 12 against Cʹ terminal truncated HasA, as reported in the Supplementary Table S2. The plate was coated with the protein of interest at 4 °C overnight, washed with 200 µl of SBB three times, blocked with 200 µl of 3% BSA for 1.5 hours at 4 °C and washed again with 200 µl of SBB three times. DNA aptamers were folded as described above in the *SELEX on magnetic beads* protocol and incubated first with an empty well as counter selection step and then with the coated protein, as indicated in Supplementary Table S2. As reported above, washing step was performed with 200 µl of SWB. The modality of the wash and the SWB composition vary according to the applied stringency of the washing conditions (Supplementary Table S2). DNA aptamers were eluted from beads-protein complex in 100 µl of nuclease-free H_2_O by heating at 95 °C, twice. PCR amplification of the recovered aptamer pool and regeneration of the ssDNA conformation were performed as described in the *SELEX on magnetic beads* protocol.

### Monitoring of SELEX progression

DNA library as well as DNA aptamer pools were amplified by qPCR with Luna® Universal qPCR Master Mix (New England Biolab, M3003X) and with the following forward (5ʹ-CGCGACGGATCTCGTTGCAT-3ʹ) and reverse (5ʹ-GGCTCCTAGCTTATCGGTCC-3ʹ) primers at 0.1 µM. The amplification protocol was 95 °C for 60 sec and (95 °C for 15 sec, 60 °C for 30 sec) x 30 cycles. After amplification, melt curve was applied: 95 °C for 60 sec, 65 °C for 60 sec and 65 °C to 95 °C (ΔT= 0.5 °C for 10 sec). Melting temperature peaks were determined by Biorad CFX Manager 2.0 software by calculating the negative first derivative of fluorescence with respect to the temperature (-*d*(RFU)/*d*T).

### Next Generation Sequencing (NGS) and secondary structure prediction

DNA aptamer pool after round 12 was amplified by PCR with Q5® High-Fidelity DNA Polymerase (New England Biolab, M0491L) and with the following forward (5ʹ-AAGTATGAATGTATGATAGTTAGTTAGTTACGCGACGGATCTCGTTGCAT-3ʹ) and reverse (5ʹ-AAGATTGATTCTTACAATGATAGTATCTAAGGCTCCTAGCTTATCGGTCC-3ʹ) primers at 0.5 µM, to adapt the sample to Illumina NGS length requirements of 150-500 bp (Amplicon-EZ, GENEWIZ). The amplification protocol was 98 °C for 30 sec, (98 °C for 10 sec, 68 °C for 15 sec, 72 °C for 20 sec) x 24 cycles and 72 °C for 2 min. PCR product was purified by QIAquick PCR Purification Kit and quantified by UV adsorption set at 260 nm. Illumina NGS of the PCR product generated paired-end data, delivered as two compressed FASTQ files, R1 (forward read) and R2 (reverse read). AptaSUITE software was used to analyze R1 and R2 reads and create a cluster table with the most enriched aptamers by identifying, counting, and clustering sequence reads^42^. Mfold web server was used to predict the secondary structure of the aptamers by setting the temperature to 25 °C and ionic conditions to 150 mM NaCl and 2 mM MgCl_2_^43^.

### Enzyme-Linked Oligonucleotide Assay (ELONA) and qPCR read-out

DNA library together with candidate aptamers HasA_1, HasA_2, HasA_19, HasA_91 and HasA_88 were evaluated by ELONA for binding to full length HasA, Cʹ terminal truncated HasA and BSA (control), in four biological replicates and according to the methodology described by Moreno and colleagues^44^. One µg of proteins was coated in triplicate on Nunc MaxiSorp plate at 4 °C overnight, washed with EBB (50 mM sodium phosphate pH 6.5, 150 mM NaCl) three times, blocked with 3% BSA for 1.5 hours at 4 °C and washed again with EBB three times. One pmol of DNA library and of each aptamer was refolded as described above in the *SELEX on magnetic beads* protocol, incubated with the proteins for 30 min at room temperature and washed ten times using 200 µl EWB (50 mM sodium phosphate pH 6.5, 150 mM NaCl, 2 mM MgCl_2_, 50 µg/ml hsDNA). Elution of both DNA library and aptamers was performed by adding 100 µl of nuclease-free water at 95 °C, heat for 5 min more at 95 °C and recover the supernatant. Each recovered sample was adjusted to the same volume of 120 µl. Recovered DNA library and aptamers were amplified by qPCR in triplicate and retrieved Ct values (average of the 9 values obtained from each biological replicate) were interpolated within the corresponding, constructed calibration curves (Supplementary Figure S3) to quantify the amount of DNA library and aptamers bound to each protein (Supplementary Table S4). Calibration standard curves were built in the range of 10^-3^ and 10^-7^ pmol (Supplementary Figure S3). Amplification by qPCR of the DNA library and aptamer candidates was performed as described in *Monitoring of SELEX progression* protocol. Analysis of statistical significance was conducted using a 2-way ANOVA with Dunnett’s posthoc test for multiple comparisons, implemented in GraphPad Prism version 10.4.2.

### Biolayer Interferometry (BLI)

Binding specificity of candidate aptamers HasA_1, HasA_2, HasA_19, HasA_91 and HasA_88, DNA Library and scramble sequence (5ʹ-GGAGCTCCAGACTGTGGCACCGACTGACTGACTGACTGAGCGACTGACTGACTGACTGGCCCAAGAGACGAGCACCCCGG-3ʹ) toward Cʹ terminal truncated HasA protein was assessed by BLI (Octet RED96 by FortéBio/Sartorius) in three biological replicates. All the DNA samples were biotinylated at the 5ʹ position to be immobilized on the streptavidin-coated biosensor tip surfaces. The kinetic binding assay was performed with 12.5 nM of DNA samples and with Cʹ terminal truncated HasA at 0, 2.5, 5 and 10 µM. The experiment settings were Baseline (60 sec), Loading (120 sec), Baseline2 (100 sec), Association (300 sec), and Dissociation (600 sec). Data analysis was performed through the Data Analysis software 11.0, raw data were subtracted by double-reference (empty biosensor and reference well), then subtracted data were fitted globally with the Rmax unlinked by sensor and with a 2:1 heterogenous model (Supplementary Table S5).

