## Supplementary Information for "DNA aptamers to interfere with HasA, the hemophore of the heme assimilation system in *Pseudomonas aeruginosa,* as a potential antimicrobial strategy"

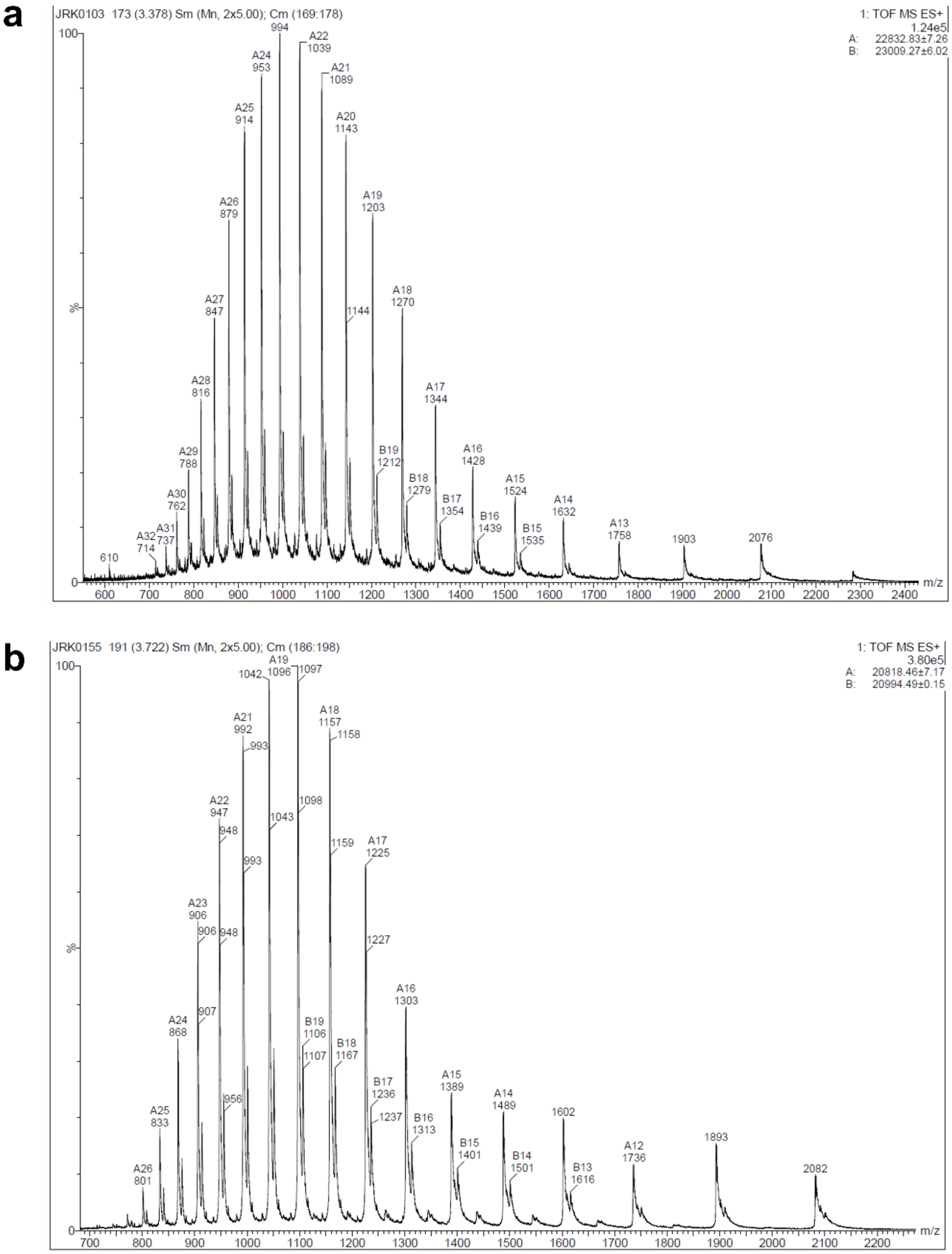


**Supplementary Figure S1.** Electrospray ionisation mass spectrometry of full length HasA and C' terminal truncated HasA. **(a)** Full length HasA shows a prominent peak at the expected size of 22.8 kDa and a secondary peak at 23.0 kDa. **(b)** C' terminal truncated HasA also shows a prominent peak at expected size of 20.8 kDa and a secondary peak at 21.0 kDa.

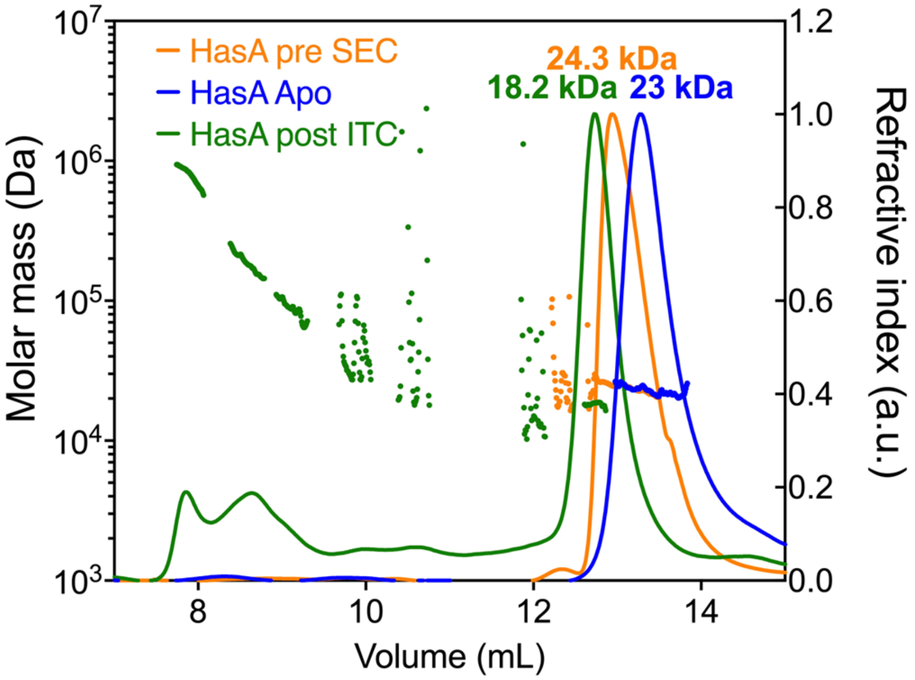


**Supplementary Figure S2.** SEC-MALS analysis of C' terminal truncated HasA in different forms: apo state (blue trace), pre SEC - holo state (orange trace) and post ITC - holo state (green trace). The assessed size of the apo state was 23 kDa, while the protein size of the pre SEC - holo state increased to 24.3 kDa, confirming heme binding. The calculated size of the protein post ITC - holo state decreased to 18.2 kDa, suggesting an issue with the analysis probably due to the low protein concentration down to a level where the mass calculation becomes less accurate. Similarly, the masses of higher molecular weight oligomers cannot be determined with any certainty.

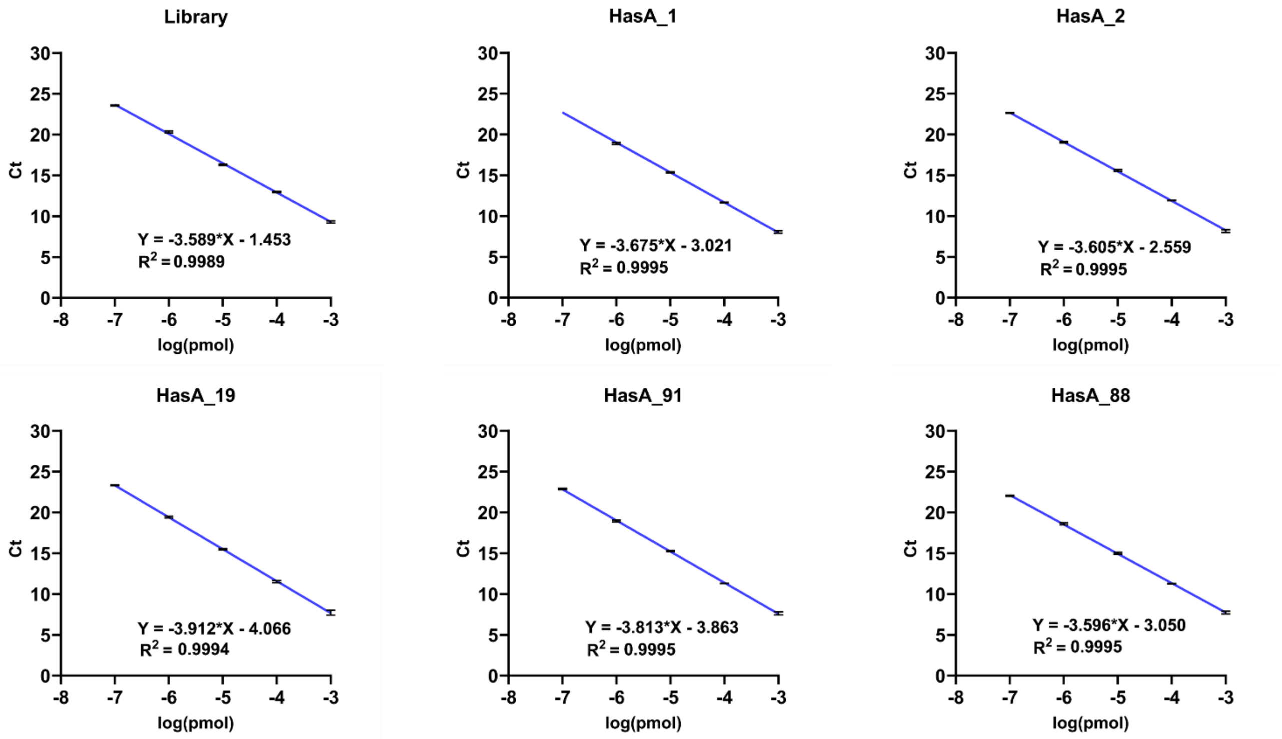


**Supplementary Figure S3.** Standard calibration curves used for DNA quantification in ELONA coupled with qPCR readout and calculated for DNA library and DNA aptamers HasA_1, HasA_2, HasA_19, HasA_91 and HasA_88 within the range of $10^{-3}$ and $10^{-7}$ pmol. Calibration curves were performed in three technical replicates showing average Ct ± SD. Curve-fitting equations and $R^2$ are reported.

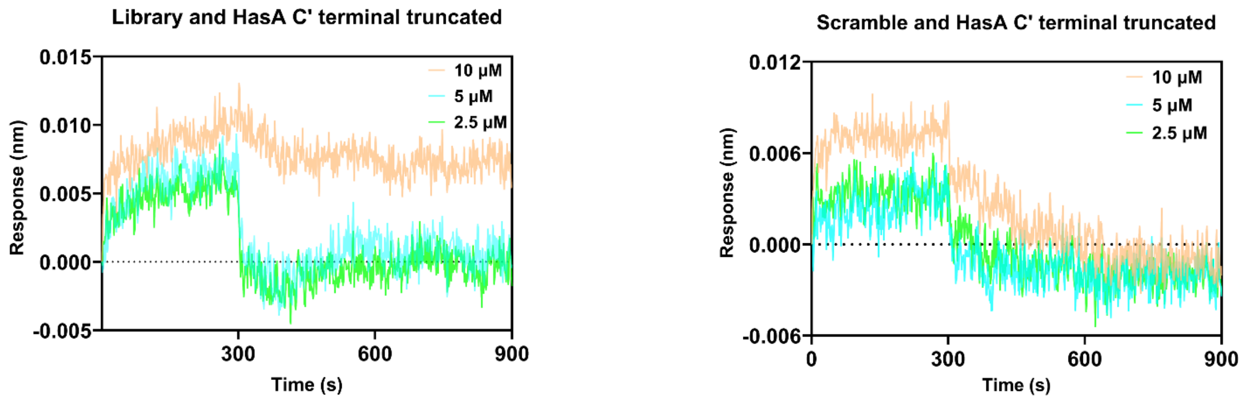


**Supplementary Figure S4.** BLI was performed in three biological replicates, binding curves from the controls (DNA library and scramble sequence) were obtained using C' terminal truncated HasA protein at concentrations of 10, 5, and 2.5 μM (orange, cyan and green curves, respectively). No significant binding responses were detected for both controls.

| Cell (M) | Syringe (M) | N (sites) | KD (M) | ∆H (Kcal/mol) | Offset (Kcal/mol) | ∆G (Kcal/mol) | -T∆S (Kcal/mol) |
|---|---|---|---|---|---|---|---|
| 19.0e-6 | 600e-6 | 1.16 ± 8.1e-3 | 113e-9 ± 29.8e-9 | -7.71 ± 0.109 | -2.18 ± 3.4e-2 | -9.48 | -1.77 |

**Supplementary Table S1.** Binding parameters from ITC assay of C' terminal truncated HasA. Cell, Syringe, N, $K_D$, ∆H, Offset, ∆G, and -T∆S are reported.

| | **R1** | **R2** | **R3** | **R4** | **R5** | **R6** | **R7** | **R8** | **R9** | **R10** | **R11** | **R12** |
|---|---|---|---|---|---|---|---|---|---|---|---|---|
| **DNA POOL** | 1 nmol | 94 pmol | 40 pmol | 30 pmol | 20 pmol | 15 pmol | 10 pmol | 10 pmol | 10 pmol | 7 pmol | 7 pmol | 7 pmol |
| **COUNTER -TARGET** | Empty-beads, 460 µg, 30 min | Empty-beads, 460 µg, 30 min | Empty-beads, 370 µg, 30 min | 2x Empty-beads, 370 µg, 30 min | Empty-beads, 280 µg, 30 min | Empty-beads, 280 µg, 30 min | Empty-beads, 200 µg, 30 min | Empty-beads, 160 µg, 30 min | Empty-well, 30 min | 2x Empty-well, 30 min | 2x Empty-well, 30 min | 2x Empty-well, 30 min |
| **TARGET** | HasA full length, 5 µg, 1 hour | HasA full length, 5 µg, 1 hour | HasA full length, 4 µg, 45 min | HasA full length, 4 µg, 45 min | HasA full length, 3 µg, 30 min | HasA full length, 3 µg, 30 min | HasA C' term trunc, 2 µg, 20 min | HasA C' term trunc, 1.5 µg, 20 min | HasA C' term trunc, 1 µg, 15 min | HasA C' term trunc, 0.5 µg, 15 min | HasA C' term trunc, 0.5 µg, 10 min | HasA C' term trunc, 0.5 µg, 10 min |
| **WASH** | 3x<br>quick<br>SWB | 5x<br>quick<br>SWB | 6x<br>3x 5 min, 2x 10 min, 1x 20 min<br>SWB | 7x<br>2x 5 min, 3x 10 min, 2x 20 min<br>SWB [200 mM NaCl] | 8x<br>3x 10 min, 3x 20 min, 2x 30 min<br>SWB [300 mM NaCl] | 9x<br>2x 10 min, 3x 20 min, 4x 30 min<br>SWB [400 mM NaCl, 50 µg/ml BSA] | 10x<br>1x 10 min, 4x 20 min, 5x 30 min<br>SWB [500 mM NaCl, 100 µg/ml BSA] | 11x<br>5x 20 min, 6x 30 min<br>SWB [600 mM NaCl, 200 µg/ml BSA] | 11x<br>5x 20 min, 6x 30 min<br>SWB [600 mM NaCl, 400 µg/ml BSA] | 12x<br>4x 20 min, 8x 30 min<br>SWB [600 mM NaCl, 400 µg/ml BSA, 50 µg/ml hsDNA] | 13x<br>3x 20 min, 10x 30 min<br>SWB [600 mM NaCl, 400 µg/ml BSA, 50 µg/ml hsDNA] | 14x<br>6x 20 min, 8x 30 min<br>SWB [600 mM NaCl, 400 µg/ml BSA, 50 µg/ml hsDNA] |
| **PCR CYCLES** | 15 | 12 | 15 | 13 | 14 | 19 | 19 | 19 | 22 | 24 | 22 | 24 |

**Supplementary Table S2.** Schematic of the aptamer selection steps from R1 to R12. DNA amount, counter-target, target, wash, and optimum number of PCR cycles for each round are reported.

| Seq Id | Aptamer pool | Enrichment (%) | ELONA & BLI |
|---|---|---|---|
| 1 | CGCGACGGATCTCGTTGCAT**GACCTCGGCGACAGACCGACCCTCGAGGCATACATGGGCA**GGACCGATAAGCTAGGAGCC | 74.600 | V |
| 2 | CGCGACGGATCTCGTTGCAT**GACCTCGGCGACAGACCGACCCTCGAGGCAAACATGGGCA**GGACCGATAAGCTAGGAGCC | 7.420 | V |
| 19 | CGCGACGGATCTCGTTGCAT**TTTGAGTGATATGATGTCGGGGGGAATAGGTTAACCGGGG**GGACCGATAAGCTAGGAGCC | 1.310 | V |
| 15 | CGCGACGGATCTCGTTGCAT**GACCTCGGCGACAGACCGACCCTCGAGGCACACATGGGCA**GGACCGATAAGCTAGGAGCC | 0.773 | |
| 16 | CGCGACGGATCTCGTTGCAT**GACCTCGGCGACAGACCGACCCTTGAGGCATACATGGGCA**GGACCGATAAGCTAGGAGCC | 0.559 | |
| 3 | CGCGACGGATCTCGTTGCAT**GACCTCGGCGATAGACCGACCCTCGAGGCATACATGGGCA**GGACCGATAAGCTAGGAGCC | 0.459 | |
| 18 | CGCGACGGATCTCGTTGCAT**GACCTCGGCGACAGACCGACCCTTGAGGCAGACATGGGCA**GGACCGATAAGCTAGGAGCC | 0.398 | |
| 9 | CGCGACGGATCTCGTTGCAT**GACCTCGGCGACAGACCGACCCACGAGGCATACATGGGCA**GGACCGATAAGCTAGGAGCC | 0.363 | |
| 70 | CGCGACGGATCTCGTTTCAT**GACCTCGGCGACAGACCGACCCTCGAGGCATACATGGGCA**GGACCGATAAGCTAGGAGCC | 0.227 | |
| 33 | CGCGACGGATCTCGTTGCAT**TTTGAGTGATATGATGCCGGGGGGAATAGGTTAACCGGGG**GGACCGATAAGCTAGGAGCC | 0.210 | |
| 5 | CGCGACGGATCTCGTTGCAT**GACCTCGGCGACAGACCGACCCTCTAGGCATACATGGGCA**GGACCGATAAGCTAGGAGCC | 0.192 | |
| 23 | CGCGACGGATCTCGTTGCAT**GACCTCGGCGACATACCGACCCTCGAGGCATACATGGGCA**GGACCGATAAGCTAGGAGCC | 0.179 | |
| 77 | CGCGACGGATCTCGTTGCAT**GACCTCGGCTACAGACCGACCCTCGAGGCATACATGGGCA**GGACCGATAAGCTAGGAGCC | 0.162 | |
| 91 | CGCGACGGATCTCGTTGCAT**CTCAAGGACACCGGACCACAGAACTGGGACACGTCACGTG**GGACCGATAAGCTAGGAGCC | 0.157 | V |
| 127 | CTCGACGGATCTCGTTGCAT**GACCTCGGCGACAGACCGACCCTCGAGGCATACATGGGCA**GGACCGATAAGCTAGGAGCC | 0.135 | |
| 17 | CGCGACGGATCTCGTTGCAT**GACCTCGGCGACAGACCGACCCTCGAGGCATACATGGGCA**GGAACGATAAGCTAGGAGCC | 0.105 | |
| 124 | CGCGACGGATCTCGTTGCAT**GACCTCGTCGACAGACCGACCCTCGAGGCATACATGGGCA**GGACCGATAAGCTAGGAGCC | 0.100 | |
| 98 | CGCGACGGATCTCGTTGCAT**GACCTCGGCGATAGACCGACCCTCGAGGCAAACATGGGCA**GGACCGATAAGCTAGGAGCC | 0.100 | |
| 88 | CGCGACGGATCTCGTTGCAT**GACCTCGGCGACAGACCGACCCCCGAGGCAAACATGGGCA**GGACCGATAAGCTAGGAGCC | 0.096 | V |
| 196 | CGCGACGGATCTCGTTGCAT**GACCTCGGCGACAGACCGACCCTCGAGGCATACATGGGCA**GGACCGATAAGCTAGGAGAC | 0.096 | |

**Supplementary Table S3.** Representation of the top 20 most abundant DNA aptamers after NGS sequencing of R12. Random regions are highlighted in bold. DNA aptamers analyzed by ELONA and BLI are marked (V).

### ELONA quantification

| pmol | C' terminal truncated HasA | Full lenght HasA | BSA |
|---|---|---|---|
| **Library** | 2.20E-05 | 9.98E-06 | 2.52E-05 |
| **HasA_1** | 7.70E-05 | 3.88E-05 | 1.05E-05 |
| **HasA_2** | 5.28E-05 | 3.62E-05 | 1.07E-05 |
| **HasA_19** | 8.58E-06 | 5.88E-06 | 9.59E-06 |
| **HasA_91** | 8.96E-06 | 4.16E-06 | 1.16E-05 |
| **HasA_88** | 1.97E-05 | 2.31E-05 | 4.51E-06 |

**Supplementary Table S4.** Quantification of DNA library and DNA aptamer quantities from ELONA coupled with qPCR readout assay. Quantities are expressed in pmol.

| Aptamer | KD (M) | KD2 (M) | KD Error | KD2 Error | kon (1/Ms) | kon2 (1/Ms) | kon Error | kon2 Error | kdis (1/s) | kdis2 (1/s) | kdis Error | kdis2 Error | Full X^2 | Full R^2 | KD (%) | KD2 (%) |
|---|---|---|---|---|---|---|---|---|---|---|---|---|---|---|---|---|
| **HasA_1** | 5.90E-07 | 2.08E-06 | 1.16E-08 | 7.93E-08 | 7.88E+02 | 9.59E+03 | 1.02E+01 | 3.18E+02 | 4.63E-04 | 1.90E-02 | 6.81E-06 | 3.36E-04 | 0.0141 | 0.9952 | 72.8 | 27.2 |
| **HasA_2** | 3.20E-07 | 2.01E-06 | 1.22E-08 | 7.60E-08 | 7.80E+02 | 7.70E+03 | 1.24E+01 | 2.59E+02 | 2.54E-04 | 1.46E-02 | 8.69E-06 | 2.76E-04 | 0.0126 | 0.9936 | 69.7 | 30.3 |

**Supplementary Table S5.** Binding parameters from BLI assay of HasA_1 and HasA_2 aptamers. $K_D$, $K_{D2}$, $K_{on}$, $K_{on2}$, $K_{dis}$, $K_{dis2}$, $X^2$ and $R^2$ are reported.
